# The first AI-based mobile application for antibiotic resistance testing

**DOI:** 10.1101/2020.07.23.216929

**Authors:** Marco Pascucci, Guilhem Royer, Jakub Adamek, David Aristizabal, Laetitia Blanche, Amine Bezzarga, Guillaume Boniface-Chang, Alex Brunner, Christian Curel, Gabriel Dulac-Arnold, Nada Malou, Clara Nordon, Vincent Runge, Franck Samson, Ellen Sebastian, Dena Soukieh, Jean-Philippe Vert, Christophe Ambroise, Mohammed-Amin Madoui

**Author notes:** authors with equal contributions (Marco Pascucci, Guilhem Royer).

## Abstract

Antimicrobial resistance is a major global health threat and its development is promoted by antibiotic misuse. While disk diffusion antibiotic susceptibility testing (AST, also called antibiogram) is broadly used to test for antibiotic resistance in bacterial infections, it faces strong criticism because of inter-operator variability and the complexity of interpretative reading. Automatic reading systems address these issues, but are not always adapted or available to resource-limited settings. We present the first artificial intelligence (AI)-based, offline smartphone application for antibiogram analysis. The application captures images with the phone’s camera, and the user is guided throughout the analysis on the same device by a user-friendly graphical interface. An embedded expert system validates the coherence of the antibiogram data and provides interpreted results. The fully automatic measurement procedure of our application’s reading system achieves an overall agreement of 90% on susceptibility categorization against a hospital-standard automatic system and 98% against manual measurement (gold standard), with reduced inter-operator variability. The application’s performance showed that the automatic reading of antibiotic resistance testing is entirely feasible on a smartphone. Moreover our application is suited for resource-limited settings, and therefore has the potential to significantly increase patients’ access to AST worldwide.

## 1 Introduction

The development of new antimicrobial agents is currently outpaced by the emergence of new antimicrobial resistance^1^ (AMR). The appearance and diffusion of AMR has become a serious health threat^2^, whose magnitude is not yet fully understood because of the lack of data, especially in areas where the access to antimicrobial susceptibility testing is difficult. A high-profile review^3^ forecasts ten million deaths worldwide by 2050. Although these numbers have been criticized^2^, these studies underline the critical health burden of AMR and the need for global data^2,4^.

Testing the susceptibility of bacteria is important for patient treatment and, if done systematically, gathering data can provide precious epidemiological information. Different test methods^5^ exist. Arguably the most widely used is the Kirby-Bauer disk diffusion test.

In this test, cellulose disks (pellets) containing antibiotics at a given concentration are placed in a Petri dish with an agar-based growth medium previously inoculated with bacteria. While the plate is left to incubate, the antibiotic diffuses from the pellet into the agar. The antibiotic concentration is highest near a pellet and decreases radially as the distance from the disk increases^6^. The bacteria cannot grow around those disks that contain antibiotics to which they are susceptible. The growth of the bacterial colony stops at a distance from the pellet which corresponds to a critical antibiotic concentration, forming a visible bacteria-free area around the cellulose disk. This is called a *zone of inhibition*. After incubation, the diameter of the zone of inhibition around each antibiotic disk is measured: the categorization of the microorganisms as susceptible (S) Intermediate (I) or Resistant (R) is obtained by comparison of the diameter against standard breakpoints^7^ established by international committees such as the European Committee on Anitimicrobial Susceptibility Testing (EUCAST) or the Clinical and Laboratory Standards Institute (CLSI)^8^.

The disk diffusion method is relatively simple, can be performed entirely by hand, requires no advanced hardware, and has a low cost. However, it is criticized for several reasons. First, it is labor-intensive and time-consuming. Second, it is subject to important inter-operator variability: accurate performance of disk diffusion testing relies on proficient technicians, starting with the quality of plate preparation (e.g. inoculum, purity)^9^. The diameter of the inhibition zone is measured by eye with a caliper or ruler and approximated to the closest millimeter^10^. However, the inhibition zone might not be a perfect disk (e.g. if the inhibition zone overlap) or if the pellet is too close to the border of the dish (see below, Figure 6a, f and h). In this case the problem of measuring a diameter is ill-posed and, together with intrinsic measurement error, introduces subjectivity and inter-operator variability in the measurement. Third, it requires an advanced level of expertise for interpretation. Sometimes, the inhibition zone diameter is not sufficient by itself to determine the susceptibility. Indeed, several mechanisms of resistance are expressed at a low level in *vitro* but have major impact in vivo and can lead to treatment failure. Moreover, susceptibility to a whole class of antibiotics or a given molecule class can sometimes be inferred from the susceptibility to another one, thus reducing the number of required tests. In those cases, interpretative reading is needed. Interpretation is based on expert rules published and updated by scientific societies, such as EUCAST in Europe^11^.

Automatic reading systems have been introduced to alleviate the drawbacks of disk diffusion AST ^12^,^13^. These systems acquire pictures of the plate and automatically measure the diameters of the inhibition zones. Most of them include an *expert system* which can elaborate interpreted results. It helps mitigate the risk that the laboratory reports erroneous susceptibility results and ensures compliance with regulatory guidelines. Commercial devices^14–16^ that automatically read antibiograms are commonly used in hospitals and laboratories, but the procedures they use are not fully disclosed. These systems aim towards great and flawless automation and a high degree of standardization of the culturing procedures in order to concurrently increase quality and turnaround times. These needs are not the same in resources-limited hospitals, where AST might be not implemented at all.

Because of their price, and hardware and infrastructure requirements, these systems are not suited to resource-limited settings such as dispensaries or hospitals in resource-limited settings. Affordable solutions are few. Image processing algorithms for automatic measuring inhibition diameters have been published ^17–21^. Among these, only AntibiogramJ^21^ presents a fully functional user-friendly software, but it operates on a desktop computer onto which the images need to be previously transferred. We believe that reducing hardware requirements to just a smartphone is key for the adoption and diffusion of such a tool. Moreover, smartphone applications are easy to adopt and use if they follow established design patterns, and they benefit from an ecosystem which facilitates setup and updates.

### 1.1 An all-in-one smartphone app for AST reading

This paper introduces the first fully offline mobile application^I^ (the App hereafter) capable of analyzing disk diffusion ASTs and yielding interpreted results, operating entirely on a smartphone. The need of such an application was identified by Médecins Sans Frontières (MSF), who often operates in low and middle income countries (LMIC) where AST is difficult or impossible to implement. The MSF Foundation brought together the people and skills needed for this application to be developed, truly believing that the App can have a great impact on the fields where MSF operates and the global fight against AMR.

The App combines original algorithms, using machine learning (ML) and image processing, with a rule-based expert system, for automatic AST analysis (see Figure 1). It embeds a clinically tested third-party expert system^13,15^ which could compensate for a lack of microbiology expertise. The user is guided throughout the whole analysis and can interact at any step with the user-friendly graphical interface of the application to verify and possibly correct the automatic measurements if needed. The whole analysis takes place on the same smartphone used to acquire the picture of the AST. Since it does not require any hardware other than a basic Android smartphone, and because it works completely offline (without internet connection), the App is suited for resourcelimited settings. Therefore, the App could help fill the digital gap, increase patients’ access to AST worldwide and possibly facilitate the collection of epidemiological data on antimicrobial resistances, the lack of which is recognized today as a major health danger^2,4^.

**Figure 1:**
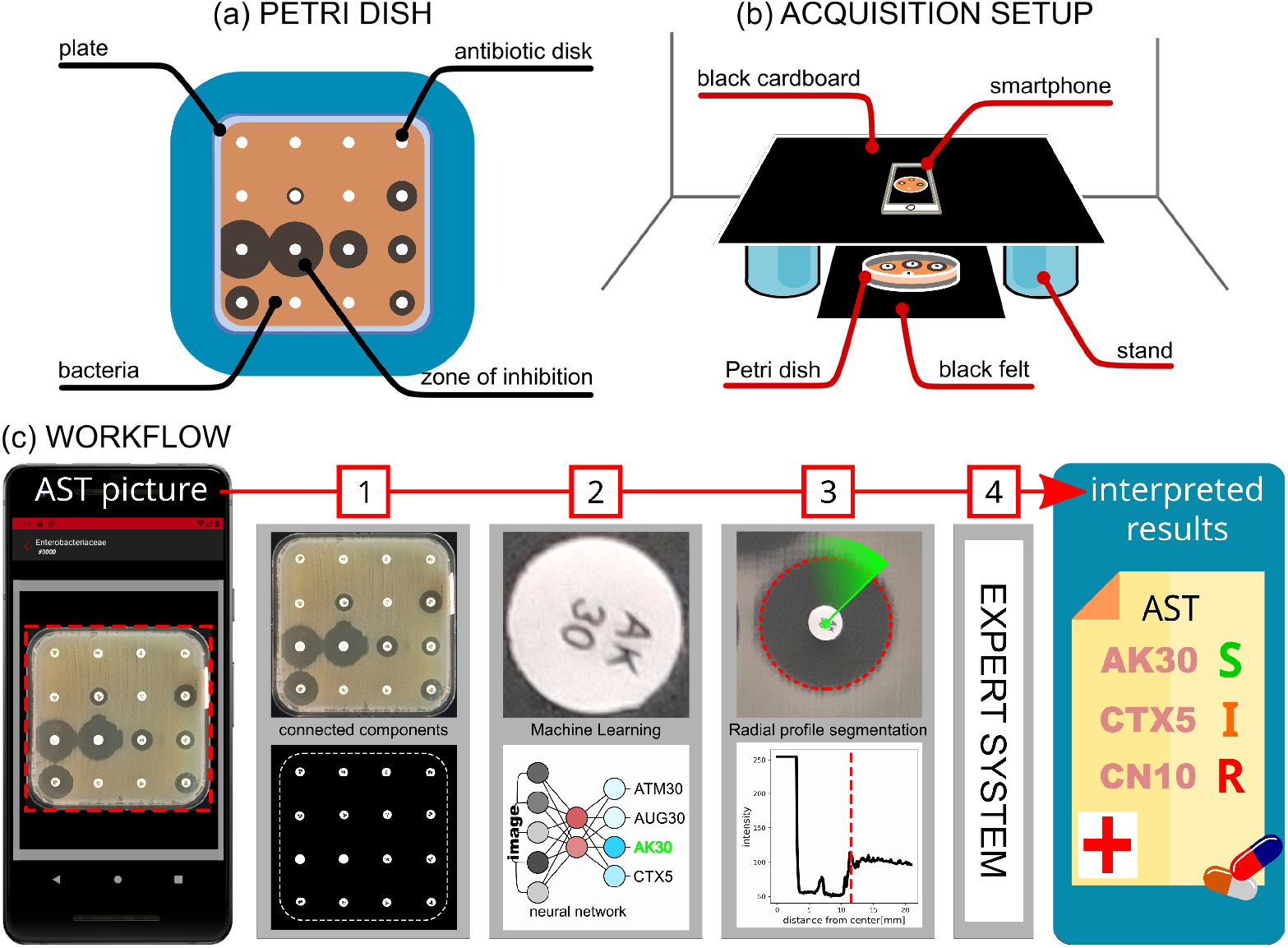
Analysis of an AST plate with the App. A prepared and incubated Petri dish (a) is positioned in a simple image acquisition setup made of cardboard (b) as stand we used two containers available in the laboratory. A picture of the plate is taken with a smartphone and the analysis follows the workflow described in (c): the Petri dish image is cropped and the antibiotic disks are found (c1); the image of each antibiotic disk is fed to a ML model that identifies the antibiotic (c2); the diameter of the inhibition zone is measured (c3) with an original algorithm. Finally, the Expert System uses the diameters to output interpreted results (c4).

In fact, the main aim of this application is to facilitate the adoption of the disk diffusion AST in resources-limited hospitals and laboratories where this test is not available yet. The App pursues this objective by partially alleviating the need of expert human resources, making the reading more reliable and provifing interpreted results. Therefore this application does not want to compete with high-end commercial systems, which can count on dedicated hardware. Nevertheless, in order to be reliable, it is fundamental that the App fulfills the minimum viable performance requirements, as we show in this work.

In the following, we demonstrate that the App’s performance is similar to that obtained with a commercial system and conform to manual reading (considered as the gold standard^10^). The application’s full automatic procedure is evaluated on antibiograms prepared in laboratory conditions both on standard and blood-enriched agar. Moreover, we explore the feasibility of a ML-based automatic detection of resistance mechanisms recognizable by peculiar shapes of the inhibition zones.

### 1.2 Image processing

The App presented in this paper is the first automatic AST reading system capable of running the whole analysis of a disk-diffusion antibiogram offline on mobile devices, from image acquisition to interpreted results. It helps laboratory technicians throughout the whole analysis process, suggesting measurements, results and interpretations. The App can be summarized in three major components:

- an dedicated image processing module (IP) that reads and analyzes the AST image,
- an expert system (ES) responsible for the interpretation of the data extracted by IP,
- a Graphical User Interface (GUI) that allows the execution of IP and ES on a smartphone.

The application’s IP module^II^ implements a novel algorithm for the measurement of the inhibition diameters (described in Methods) and uses ML for the identification of the antibiotic disks, which is unprecedented in this kind of applications. The IP module consists in a C++ library developed on OpenCV^22^ and Tensorflow^23^. The choice of C++ makes our library exploitable in various contexts, including desktop computers and Android and iOS mobile devices. Moreover, the library has a Python wrapper, useful for application prototyping, image batch processing and benchmarking.

The App includes an expert system capable of performing coherence checks on the raw susceptibility and providing interpreted results, with extrapolation on non-tested antibiotics and clinical commentaries. The expert system’s knowledge base is provided and regularly updated by i2a (Montpellier, France)^13^ and based on up-to-date EUCAST expert rules^11^. The expert system’s engine was completely developed in TypeScript and works completely offline within the App.

Commercial AST reading systems use built-in image acquisition devices (cameras and scanners) to ensure input consistency. The App works on images of ASTs taken directly with the phone’s camera, with no additional external acquisition hardware. This inevitably introduces a certain variability in the image quality. We tackle this issue by introducing a simple set of guidelines for image acquisition (see supplementary 7.1). These guidelines are designed to optimize image quality, and therefore to reduce the need of heavy post-processing and the risk of numerical artifacts. For the same reason, perspective distortions are not corrected. Instead, we developed a simple acquisition setup (Figure 1) which ensures parallelism between the dish and the camera’s image plane. The acquisition guidelines are conceived to be inexpensive and easy to implement and integrate in the laboratory routine. Since smartphone cameras are not designed for quantitative measurements, we provide a simple method to assess the camera’s optical distortions with a numerically generated AST image. Moreover, while taking the picture, the application uses the device’s gyroscopes and accelerometer, if available, to force the device orientation (parallel to ground, to avoid perspective distortions) and stability (to avoid motion blur). The application also displays a visual frame that helps center the Petri dish in the picture. Although Petri dishes have standard shapes (square or disk), we do not rely on this assumption for the analysis.

The IP module analyzes an AST picture in three different sequential stages: plate cropping, detection of antibiotic disks and inhibition diameter measurement. The resulting diameters are used to categorize the susceptibility of the bacteria and interpret the results. These stages are described in Methods and summarized in Figure 1. Once the antibiogram picture is taken, the Petri dish is cropped out, the antibiotic disks are found and classified according to their label, the diameter of the inhibition zones is measured and translated into susceptibility results. At this point the expert system verifies the coherence of the measurements to highlight possible errors and finally returns the interpreted results. Some screenshots of the App are shown in Figure 2 and a complete video demo is available online^III^.

**Figure 2:**
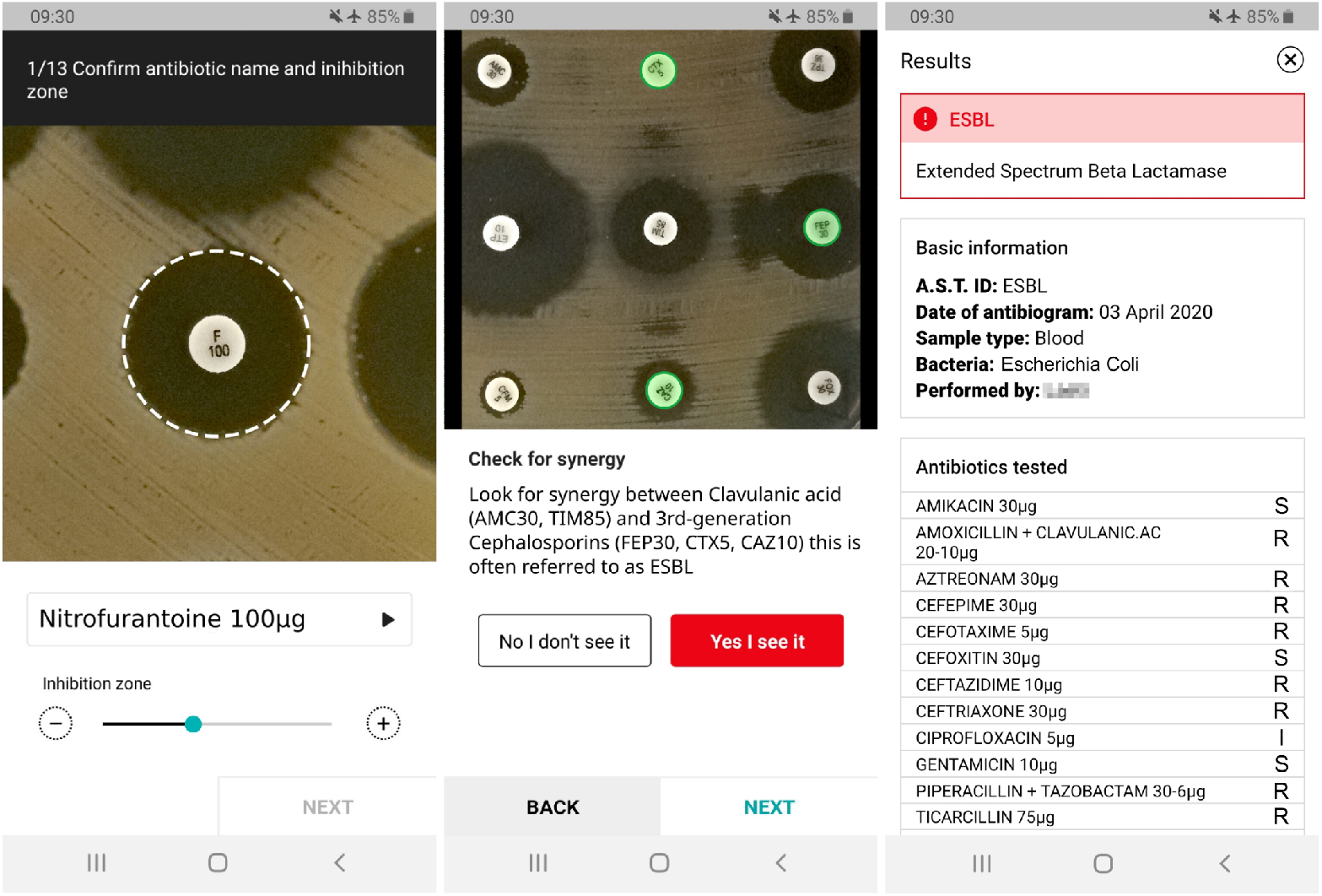
Screenshots of the App in action. Left image: the App displays a zoomed image of an inhibition zone and indicates with a dashed circle the automatically measured diameter and the detected antibiotic. The user can edit the results with the controls below the image. Central image: the application can ask the users if they see the peculiar shapes of inhibition zones associated with certain resistance mechanism. Right image: at the end of the analysis, the interpreted results are shown to the user.

### 1.3 Preparation of antibiotic susceptibility tests

In order to evaluate the App’s performances we ran the fully automatic analysis procedure (without any manual intervention or correction) on three sets of antibiograms (A1-3) described in Table 1.

**Table 1:**
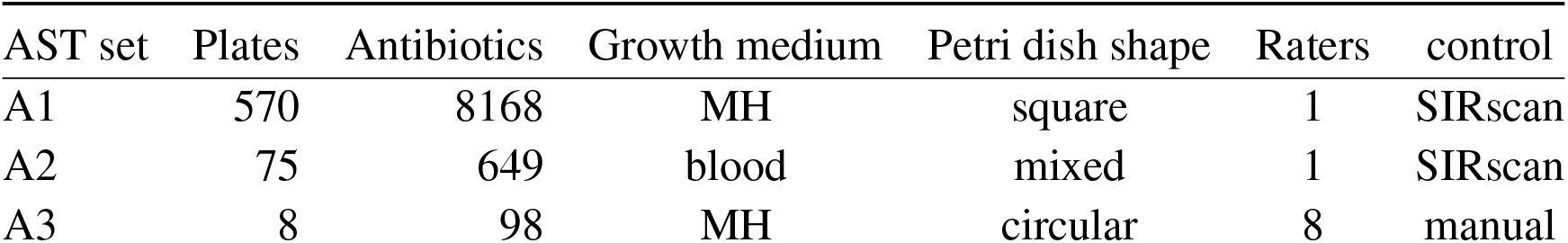
Description of the AST sets used for the performance evaluation of the automatic reading. For each data-set the columns indicate: the number of single Petri dishes in the data-set, the corresponding total number of antibiotic disks, the type of growth medium, the shape of the plates, the number of independet raters measuring the diameters, the reference used as control diameter.

| AST set | Plates | Antibiotics | Growth medium | Petri dish shape | Raters | control |
| --- | --- | --- | --- | --- | --- | --- |
| A1 | 570 | 8168 | MH | square | 1 | SIRscan |
| A2 | 75 | 649 | blood | mixed | 1 | SIRscan |
| A3 | 8 | 98 | MH | circular | 8 | manual |

AST groups A1 and A2 consist of 571 and 74 antibiograms prepared during working routine in the microbiology laboratory of the University Hospital in Créteil, France. The samples were collected from patients of the hospital and the preparation and analysis of the AST was not designed primarily for our study but followed the normal hospital procedures. AST set A3 consists of 8 Petri dishes prepared in the Hospital of Médecins Sans Frontières in Amman, Jordan. In the case of this set, the plates were inoculated with microorganisms purchased from the American Type Culture Collection (ATCC) and routinely used for quality control. Such strains are among the main pathogens and have known inhibition diameters to various antibiotics. Species distribution, preparation information and other details are reported in the Methods for all three sets.

### 1.4 Data acquisition

All plates in AST groups A1 and A2 were imaged with a smartphone camera (Honor 6x with a resolution of 12 megapixel). Since the App was still under development at the time these images were taken, we used the default Android camera application for acquisition. Then the images were analyzed with the App’s full automatic procedure (without manual intervention). As control inhibition diameters for sets A1 and A2 we collected the measurements effectuated by the laboratory technician using a commercial automatic reading system (SIRscan^13^, i2a, Montpellier, France). The control diameters measured by the technicians with the SIRscan system were extracted retrospectively from the hospital database, since these antibiograms were performed during routine analysis in the hospital. The SIRscan system allows for correction on automatic measurements. Nevertheless, for productivity reasons, the technician did not always adjust the diameters if the adjustment did not yield a different categorization result, therefore diameters can be unadjusted even if they give the right susceptibility categorization.

Among the pictures of AST groups A1 and A2, we selected *standard* and *problematic* pictures according to the following criterion: if for more than two antibiotics in the picture we found an absolute diameter difference between the App and control values of more than 20 mm, we considered the picture *problematic*, otherwise it was considered *standard*. The *problematic* images are often associated with plates with defects or show very low inhibition-to-bacteria intensity contrast (due to low bacteria pigmentation and/or low illumination conditions). Nevertheless most of the low contrast images in A1 and A2 where classified as *standard* (see Figure 3).

**Figure 3:**
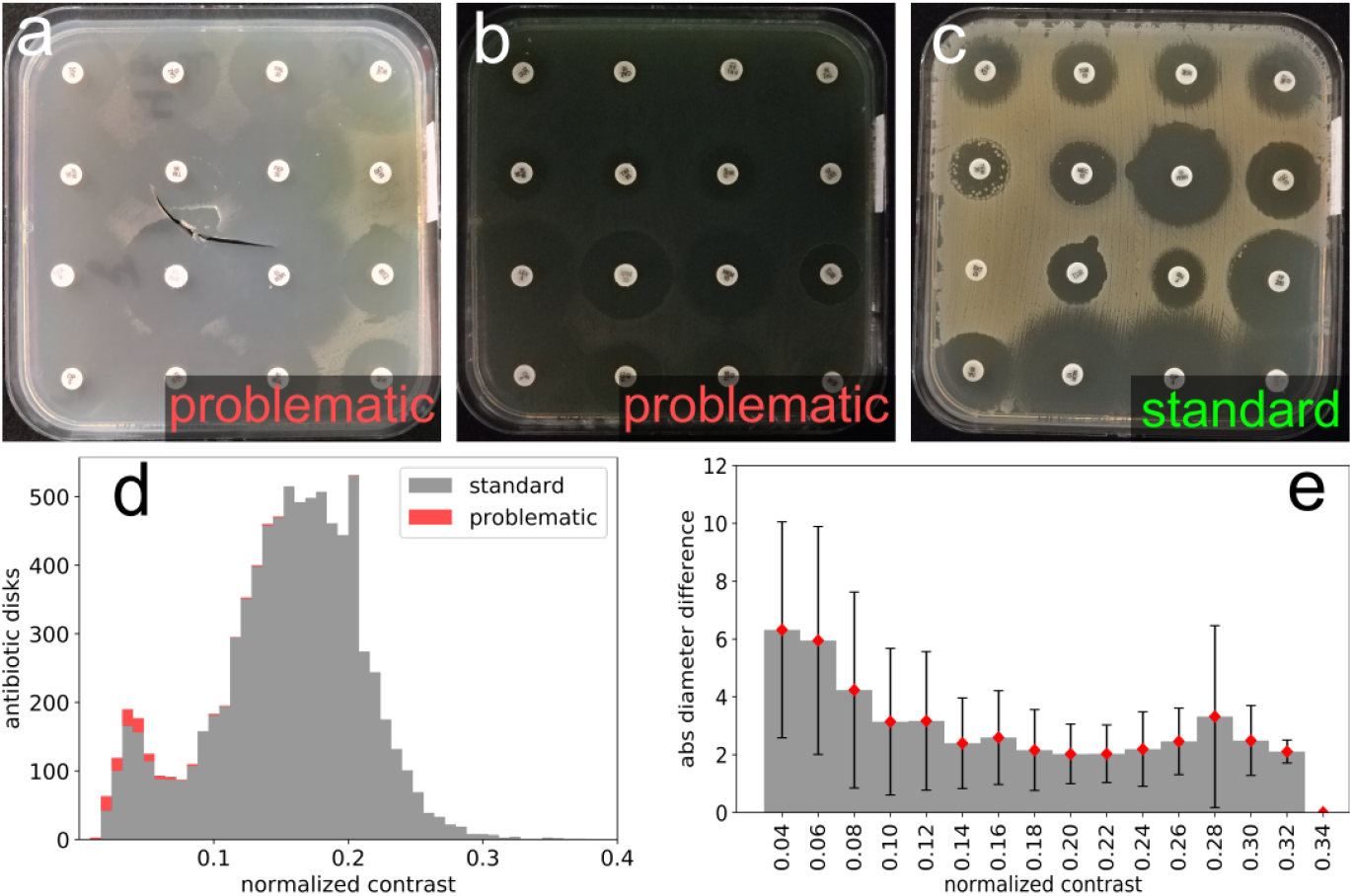
A few *problematic* images have been identified in the dataset. They correspond to damaged plates (a) and images with very poor visible contrast between the bacteria and the inhibition (b). Even by eye, it is difficult to clearly identify all the inhibition zones in these cases. For comparison, a standard image looks like (c). The coupled effect of bacteria pigmentation and variable illumination produces a considerable variability in the bacteria-to-inhibition intensity contrast (a,b,c). The histogram in (d) shows the distribution of image contrast for *standard* and *problematic* images in AST set A1 (the contrast is defined here as the difference between the central intensity level of bacteria and inhibition): problematic images (in red) are a small fraction of the total, mainly concentrated in the lower contrast region. Finally, (e) shows the observed mean diameter difference versus contrast: low contrast images yield worse results.

All plates in A3 were imaged with the App on a smartphone (Samsung A10, 12 megapixel camera) by eight different laboratory technicians to take into account inter-operator variability (e.g. plate position, contrast and random noise, see Figure 6h,i,j). The resulting 64 images were analyzed with the App’s full automatic procedure. As a control, each AST was measured manually with a ruler by the same eight lab technicians. In this way, each inhibition diameter was measured eight times.

### 1.5 Benchmark

The diameters of the inhibition zones read with the App’s automatic procedure were compared with the control diameters. For every diameter, we calculated the absolute difference with the corresponding control value. The susceptibility categorization (SIR) of the antibiotics was made for both the App’s procedure and control, by comparing the inhibition diameter of each antibiotic to the breakpoint defined in the EUCAST guidelines^7^. Antibiotics for which a breakpoint was not provided are excluded. In order to evaluate the App’s performance, we compared the susceptibility categorization of the automatic procedure to the control one. Following the same terminology proposed by ^17,20,21^, we calculated the *agreement* as the rate of identical categorization; disagreement is classified as *very-major, major* and *minor*. Very-major disagreement occurs when an antibiotic is categorized S (Susceptible) while the control is R (Resistant), major error corresponds to a categorization of R with control S and minor disagreement is any other categorization error involving the Intermediate value I (Intermediate). As a measure of agreement between the App and control, we calculated the unweighted Cohen’s kappa-index^24^.

## 2 Results

### 2.1 Image processing performance

The App’s IP procedure proved its reliability at each step. For each data-set, the photos were automatically cropped to isolate the Petri dish. The automatic crop procedure never failed if the image respected the acquisition protocol (see Supplementary 7.1). The automatic pellet detection correctly found all antibiotic pellets in A1 and A2. In A3 0.5 % of pellets where missed (false negatives). Half of the missed pellet show visible flaws (see Figure 11) and should not be considered in the analysis, according to experts advice. False positives never occurred (other objects wrongly identified as pellets). In case a pellet was missed, the users can add it with the help of the graphical interface.

**Figure 11:**
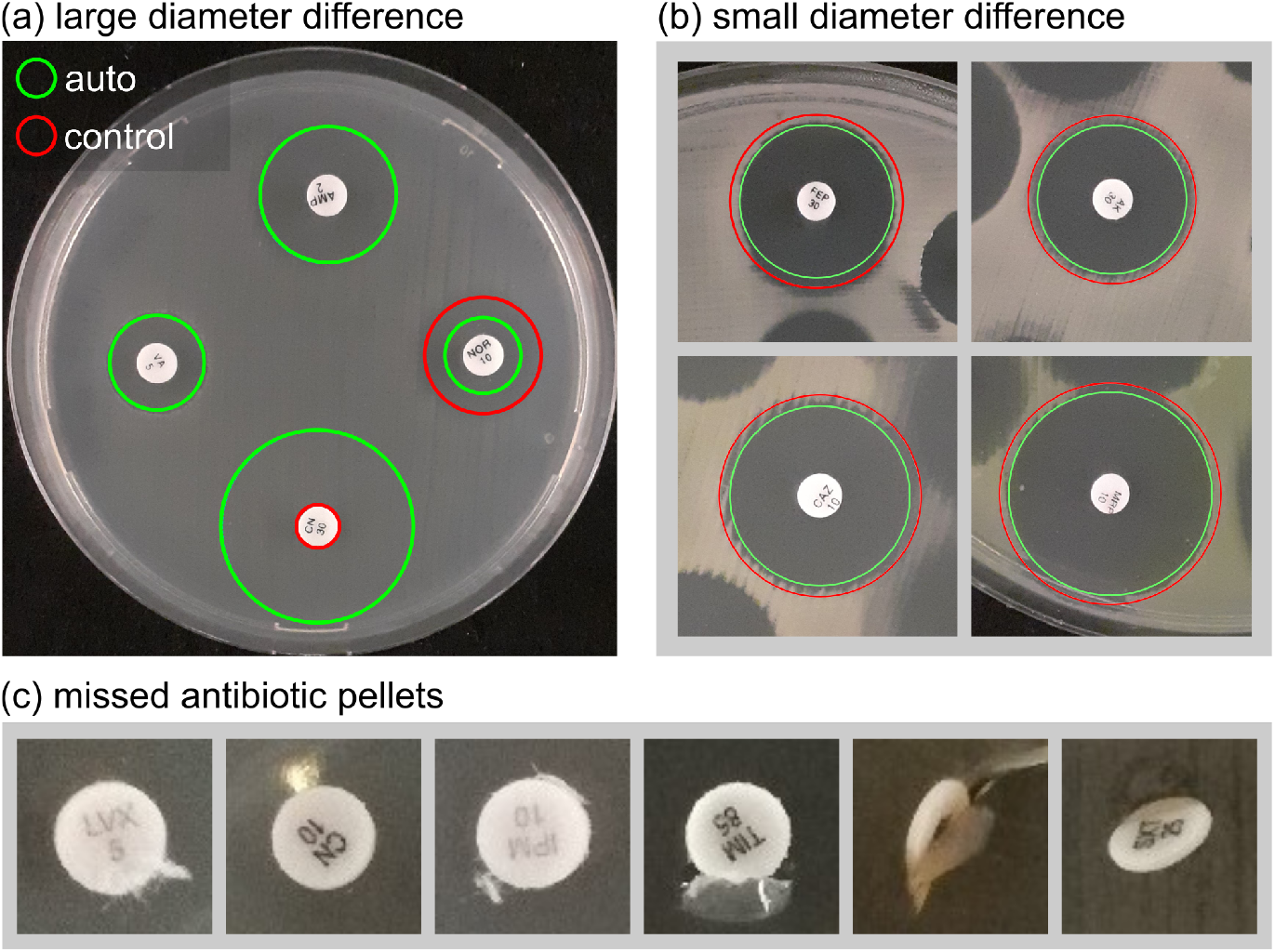
Automatic reading mistakes. Cases of large mistakes in inhibition diameters happen sometimes when the bacteria-inhibition contrast is low (a). Smaller diameter differences between the App and reference are probably due to the intrinsic measurement method: by visualizing a circle the App increases the measurement precision. Figure (c) shows some examples of pellets we were not able to automatically find. The three last pellets have broken the agar, hence should not be read anyways.

The antibiotic labels were always correctly interpreted in A3, even if the image was not perfectly focused or the text was damaged (by bad printing or by positioning it with tweezers). In data-sets A1 and A2 the accuracy was 98%. Misclassification happened in cases of very poorly printed labels or for pellets from non-supported providers, on which the ML model was not trained. To overcome this problem, in the app we calculate a confidence value for each classification yielded by the model in order to reject misclassified labels and ask the user to identify them by eye (see Supplementary 7.3 for detailed results). Moreover, since the whole process is supervised by the user, misclassifications can easily be corrected.

Furthermore, we demonstrate a proof of concept for the ML classification of resistance mechanisms. We examined two clinically-relevant resistance mechanisms that are traditionally detected by the presence of non-circular inhibition zones. By training simple convolutional neural network models on the relevant portions of AST images, we obtained encouraging results: Accuracy higher than 99.7% in detecting induction (indicative of MLSb-inducible resistant Staphylococcus aureus) and 98% in detecting synergy indicative of ESBL production (see Supplementary 7.5 and 7.6). However, due to user experience considerations in combination with concerns about model transferability, we ultimately determined not to incorporate these resistance mechanism-detecting models into the App. We do not exclude reconsidering this approach in future version of the application if we can generalize it to a larger and diverse data-set. Instead, every time a resistance mechanism can appear in a culture (given the bacteria species and the tested antibiotics) the application will systematically ask the user to verify the presence of the associated shape, showing illustrated examples (see Figure 2).

### 2.2 Susceptibility categorization

Our new diameter measurement approach yielded good classification results over most of the available images, as showed in Table 2. Only a small fraction of the pictures, classified as *problematic* in Table 2 (1.5%, 2.6% and 0 in A1, A2 and A3 respectively), produced major discrepancies. Details, metrics and a discussion on image contrast can be found in Methods, and in the supplementary material (Table 4 and Radial profiles). Overall, diameter measurements allowed a susceptibility categorization agreement of at least 90% for all three antibiogram sets (all images included). The categorization results are reported in Table 2.

**Table 2:**
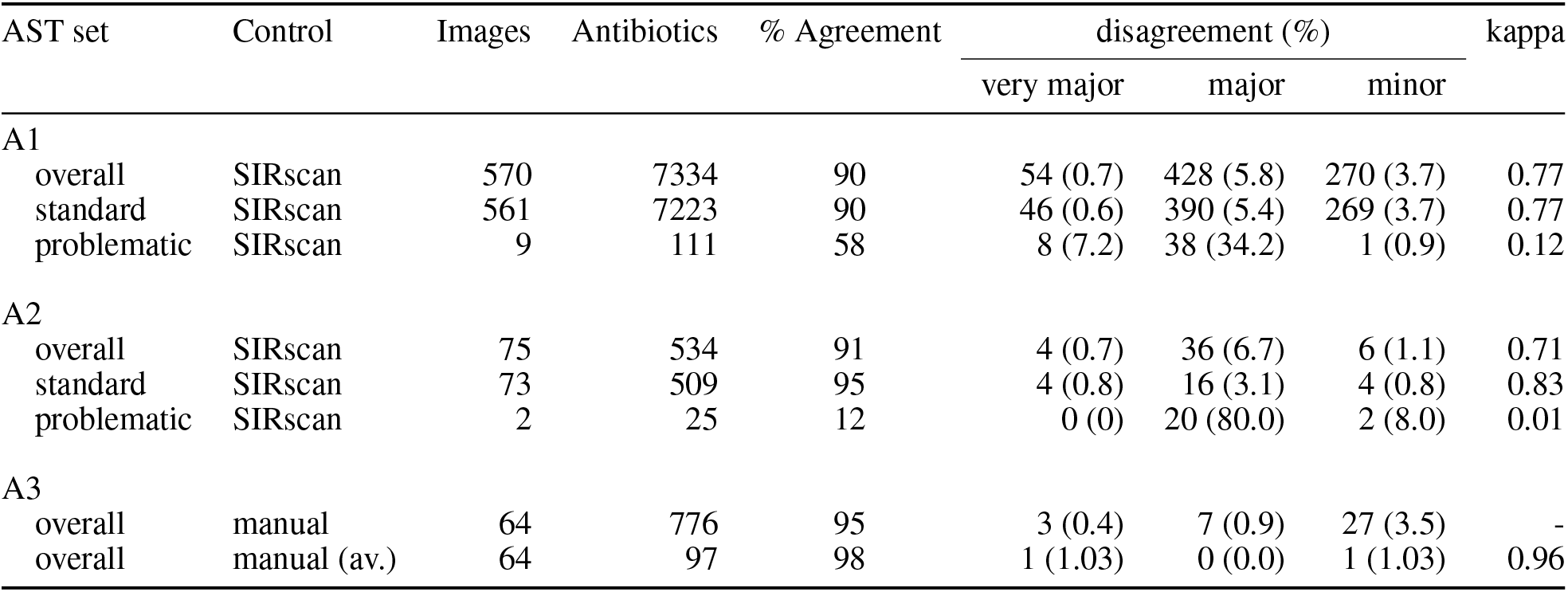
Categorization agreement between the App automatic procedure and control. The number of antibiotic reported here is the number of those for which clinical breakpoints are provided by the EUCAST^7^. The lines of this table present the agreement/disagreement for all antibiogram sets (A1, A2 and A3). For each line we specify: the control of diameter values, the number of analyzed images and corresponding antibiotic pellets, the agreement and disagreement (as defined in the text), the Cohen’s Kappa coefficient as another measure of agreement. The label “overall” means that all pictures are considered, whereas “standard” and “problematic” stand for the respective images subsets (see Benchmark section). The plates in sets A1 and A3 were grown on standard Mueller-Hinton (MH) growth medium, whereas we used blood enriched MH in A2. In the last line of the table, “av.” stands for average. In this line we used as control diameters the average value across the measurements of all eight technicians.

The actual distribution of the diameter differences among manual, automatic and assisted (corrected by the user) readings of A3 are shown in Figure 4. In general, we observe that the manual measurements (done with a ruler) are on average slightly larger than the automatic and assisted ones. The assisted measurements are done by the technicians directly on a smartphone. The graphical interface displays a circle centered on the antibiotic disk that the user can adjust in diameter until it fits the zone of inhibition. With this kind of visualization, the measurement is easier and more accurate than the one done with a segment (which is the case of the ruler). We argue that most of the diameter difference between automatic reading and control are due to the difficulty of measuring with a ruler the diameter of inhibition zones which are not perfectly circular (see Figure 11 and Figure 6). Instead, more accurate measurements are obtained by adjusting a circular guide as in the App. The positive effect of measuring with the App is also visible in Figure 4, which displays the inter-operator average diameter difference. As expected, we observe that using the App lowered inter-operator variability.

**Figure 4:**
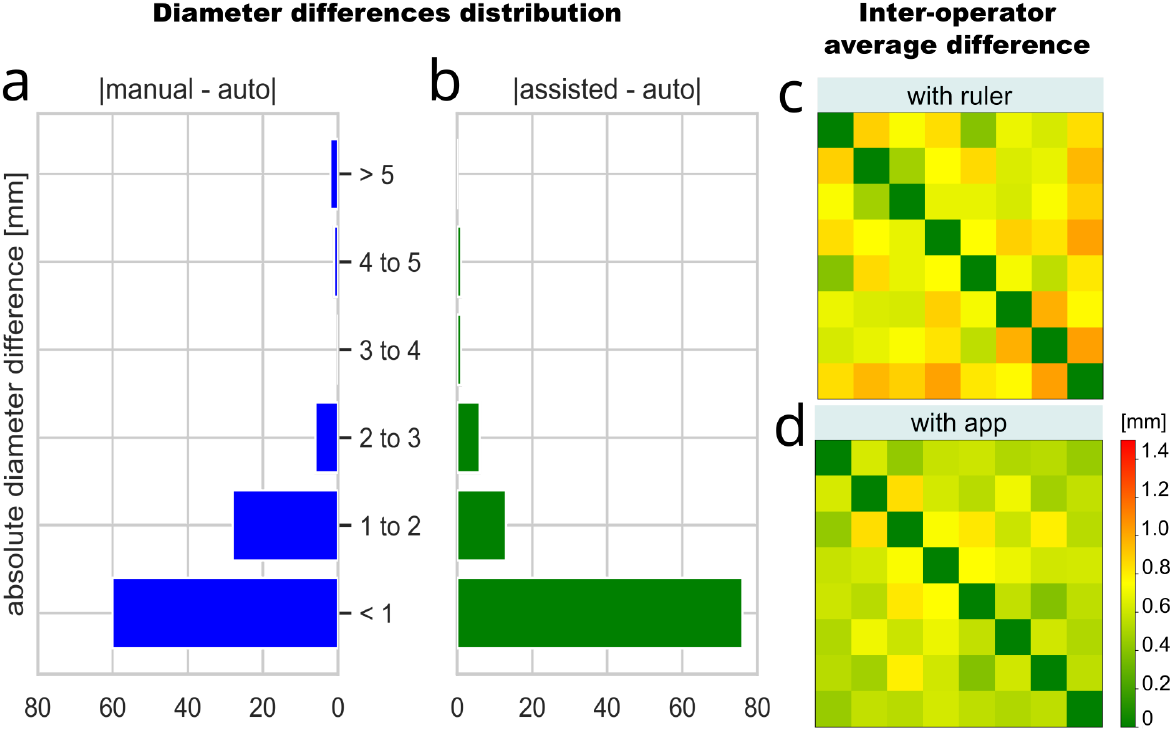
Visualization of the benchmark results on data-set A3. The histograms (a,b) show the distribution of the absolute diameter differences between the App’s automatic procedure (*auto*) and the manual measurement with ruler (a) as well as with the diameter adjusted on the smartphone by the technicians (*assisted*, b). On the right, the heat-maps show the average absolute measurement difference among the eight technicians (given 2 readers *i* and *j*, square *i*, *j* represents the difference between them) measuring with the ruler (c) and in assisted mode with the App (d). The assisted measure seems to reduce inter-operator variability.

The choices we made in building the App and the acquisition setup are made in order to facil-itate its adoption in the laboratory routine in low-resource settings. The counterpart of this choice is a certain difficulty in obtaining a constant image quality (notably because of the intrinsic variability of smartphone hardware and software). The acquisition setup and GUI assistance are there to limit this variability, but can not standardize the image acquisition at the level of commercial systems with dedicated hardware. Nevertheless the data in this study display a certain variability (especially in contrast, as shown in Figure 3) with which the application’s IP could easily cope. Strong light reflections and other important issues will result in evident wrong readings, which are easily detectable by eye with the App’s user interface. We remarked that, even without training, users can easily notice such problems and adapt the acquisition setup in order to eliminate them in future acquisitions.

Finally, we compared the App to other existing systems. The categorization agreement and errors observed in this studies among the App’s automatic procedure and control are similar to those of other systems (free and commercial) found in the literature (see Supplementary Table 5). The image treatment has been designed to perform on mobile devices, and does not require the user’s intervention to optimize image features (e.g. contrast). We obtained consistent results even by down-scaling the antibiogram pictures up to a resolution of 1 megapixel (see Supplementary Table 6). The whole reading of one antibiogram (12 megapixel picture, 16 antibiotic disks) takes less than 1s on a PC using one 2.3 GHz Intel Core i5 processor, 1.5 s on a high-end smartphone (Pixel 3 released in 2018) and 6.6 s on a low-end smartphone (Samsung A10), still much faster than manual reading.

For the matter of hardware compatibility, as of now, we have tested three smartphone models (Google Pixel 3A, Honor 6x, Samsung A10) ranging from high-to low-end. We thoroughly tested the most affordable and available model (Samsung A10) and can recommend it as a trusted device. In the future, we will maintain a list of recommended devices associated with the App.

## 3 Discussions

In this paper we have presented the first fully-offline smartphone application capable of analyzing disk-diffusion antibiograms. The App assists the user in taking a picture of an disk diffusion AST plate, measuring and categorizing zones of inhibition, and interpreting the results. The analysis is performed entirely on the same device used to acquire the picture of the antibiogram.

The App shows performances similar to other existing automatic reading systems. In particular, the automatic inhibition zone diameter reading is consistent with manual reading (gold standard). The observed accuracy is therefore considered satisfactory for usage in an AST reading system assistant. A user-friendly interface makes it easy for the user to adjust the automatic results if needed. We tested the App on antibiograms prepared with standard Mueller-Hinton (MH) growth medium as well as with MH supplemented with blood, used for fastidious organisms and obtained similar results.

We built and trained two ML-based image classification models to identify resistance mechanisms. The accuracy results are encouraging, but given the relatively small training sets, we consider the risk of over-fitting too high for the scope of such a medical device. Nevertheless, these cases are handled by the integrated Expert System, which asks the user to confirm/exclude the presence of such shapes, when likely to happen.

The App aims to encourage the implementation of disk diffusion AST in resources-limited hospitals and laboratories where antibiograms are not routinely used or poorly interpreted. It does this by simplifying the measurement task and by providing an interpretation tool, offline, on a simple smartphone with camera. The App is part of the mobile-Health^25^ (mHealth) revolution, which aims to increase patients’ access to testing, to aid in their treatment and to decrease the digital gap in the world. Our hope is that, the App could help fill the digital gap and increase patients’ access to AST worldwide.

Further clinical investigations using the App in MSF hospitals will estimate the patient benefit enabled by AI-based antibiotic resistance testing. Pending the results, the mobile application will be released and open-sourced to the public under the name of AntigbioGo. The App will support selective reporting of antibiotic sensitivity^26^, an important component of the an antibiotic stewardship strategy. It will offer the option of contributing data to global AMR surveillance with institutional bodies in place such as the WHO program GLASS ^IV^ (Global Antimicrobial Resistance Surveillance System) and/or WHOnet^V^ in order to facilitate the collection of epidemiological data on antimicrobial resistances.

## 4 Data Availability

The datasets analysed during the current study are available at this URL: http://stat.genopole.cnrs.fr/ast.zip. The data used for training the ESBL and D-SHAPE proof-of-principle models are available from The MSF Foundation upon reasonable request.

## 5 Code Availability

The image processing library described in this paper is distributed as open-source software^IV^. As of today, The App and its source code are available for research purposes upon request^VII^. We plan to release the App as open-source software after approval of the CE authority as a clinical device. Fondation Médecins sans Frontières sees the CE mark of its app solution (as a self-certified IVD SW) as a means to demonstrate and communicate on the quality and robustness of this digital tool for a non-profit. Waiting to comply to the IVDD Directive 98/79, they do not want to grant open access until they get this certification, as it could imply legal pursuit in France for the legal manufacturer (Fondation MSF) distributing a medical device without certification.

## 6 Methods

### 6.1 Detail of the AST sets

In this section we give a detailed description of the AST data-sets used for benchmark. The specific characteristics of each data set are summarized in Table 1. The bacterial species appearing in this study are reported in Figure 5.

**Figure 5:**
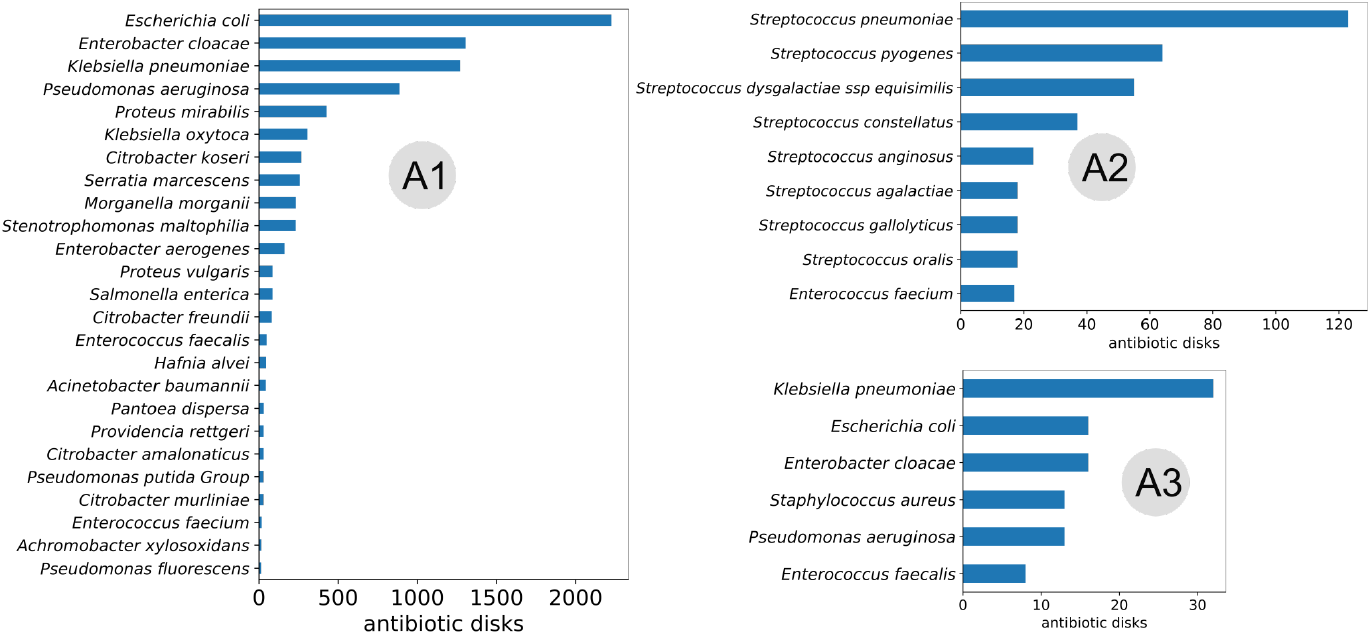
Distribution of species in the AST groups used for benchmarking.

The plates were inoculated with 0.5 McFarland of a pure culture of the studied organism. Then antibiotic disks were positioned onto the plates with a dispenser gun (A1 and A2) or by hand (A3) and the plates were incubated 16 to 24h under aerobic or 5% CO2 conditions depending on the species.

- **Data-sets A1 and A2**: More than 91% of the plates in these data-sets were square; the remaining 9 % were circular. The antibiotic disks were bought from i2a (Montpellier, France) and positioned onto the plate with a dispenser gun. Antibiograms were performed according to the EUCAST^10^ recommendations. Standard Mueller-Hinton agar was used in A1 whereas in A2 we used MH-F agar (blood-agar) for fastidious organisms (Biorad, Marnes-la-Coquette, France). The bacteriology laboratory of Créteil University Hospital is accredited under ISO15189 (Accreditation certificate N°8-3372 rév.9) therefore antibiotic disks and culture media are routinely quality checked.
- **Data-set A3**: This data-set consists of 8 Petri dishes, all circular. The antibiotic pellets were produced by Liofilchem and positioned by hand with metallic tweezers. Specifically, the Petri dishes have been inoculated with the following ATCC dried microorganisms:

1. – *Pseudomonas aeruginosa* (ATCC 27583)
2. – *Klebsiella pneumoniae* carbapenemase producer (ATCC 700603)
3. – *Klebsiella pneumoniae* SHV-18-ESBL-producer (ATCC 700603)
4. – Fluoroquinolone susceptible *Escherichia coli* reference strains (ATCC 25922)
5. – Methicillin-Resistant *Staphylococcus aureus* (NCTC 12493)
6. – Vancomycin-sensitive *Enterococcus faecalis* (ATCC 29212)
7. – Gentamicin-resistant *Enterococcus faecalis* (ATCC 49532)

### 6.2 Image processing procedure

#### 6.2.1 The custom IP library

The IP module of the App consists of a custom C++ library and it is endowed with a Python wrapper module and a quick-start documentation^VIII^. We have deliberately developed the IP as an standalone module in order to facilitate its use in other projects involving, for example, batch-processing of many images, or the integration in a Desktop application with the development of dedicated image acquisition hardware. Our aim for the App is to keep the hardware and setup as simple as possible, which is why we adopted the smartphone strategy. Nevertheless, in other projects, the mobile phone could be replaced with a small cost device like a Raspberry Pi with a camera still using our IP library.

#### 6.2.2 Plate cropping

The first raw input to IP is an AST picture, which consists of a plate (Petri dish) to be cropped out from the remaining background (see Figure 1). Cropping is accomplished with the *GrabCut* algorithm^27^, with the assumption that the plate is approximately centered in the image (i.e lies within the frame displayed on the camera screen). From the cropped plate image, we extract the dominant color, to distinguish the type of growth media (MH or blood enriched HM), and the shape of the plate (round or square). Finally, the image is converted to gray-scale for further processing.

#### 6.2.3 Detection of antibiotic disks

The image of a plate (Figure 1a) contains three main distinct components: the bacteria-free growth medium, the bacteria-covered growth medium, and the antibiotic disks. The latter are white round cellulose disks of known constant diameter (usually 6mm). The precision of the whole automatic analysis depends on the accuracy of measuring their position and diameter in the image. Since the disk radius is known as 6mm, the average disk radius in pixels is used to calculate the picture scale (pixel-to-mm ratio). Then, the inhibition diameters are measured from the disk center. The App features a fully-automatic method to measure antibiotic disks’ positions and diameters based on intensity and shape, followed by user-assisted correction. (see Supplementary 7.2)

Each pellet is printed with the acronym of the antibiotic it contains. There are only a few dozen antibiotics used in AST. Nevertheless, the acronym and the print features (font, shape, size, contrast, etc.) depend on the manufacturer of the antibiotic disks. The acronym of each antibiotic disk in the analyzed antibiogram must be read in order to retrieve the corresponding breakpoint for susceptibility categorization. Previously-proposed methods for reading these acronyms compared the image moment invariants ^20^ or used ORB (Oriented FAST and rotated BRIEF) descriptors^21,28^. For this task, we chose ML and trained a Convolutional Neural Network (CNN) model with Tensorflow^23^ (see Supplementary 7.3 for details). We trained the model on a total of 18,000 images of antibiotic disks from two different manufacturers (resulting in 65 unique labels) and achieved 99.97%. In order to limit the out-of-distribution error (wrong classifications of disks on which the model was not trained), we used an ensemble of 10 models and set a threshold on the output entropy. (see Supplementary 7.3 for details). The ML model showed to work also on poorly-printed disks and out-of-focus or low-resolution images. Interestingly we observed that even if the the printed text is damaged when the disks are placed manually (using metal tweezers), the classification is always correct.

#### 6.2.4 Inhibition disk diameter measurement

The inhibition zone diameter is the diameter of the largest circle centered on the antibiotic disk that does not include any bacteria; that is, the largest circle that can be drawn in the inhibition zone without touching any bacteria. In the easiest case, the inhibition forms a disk-shaped halo around the antibiotic pellet, but sometimes the disk is not well-defined, for example because of the overlap of several inhibition zones or because the antibiotic disk lies close to the plate borders (see Figure 6g). The bacteria-to-inhibition intensity contrast in the image depends on the bacteria species. Also, illumination can vary both among different images and within the same image. The observable effect is a visible difference of contrast in the AST images (see Figure 6h,i), especially when taken with a mobile phone where the illumination conditions cannot be controlled.

**Figure 6:**
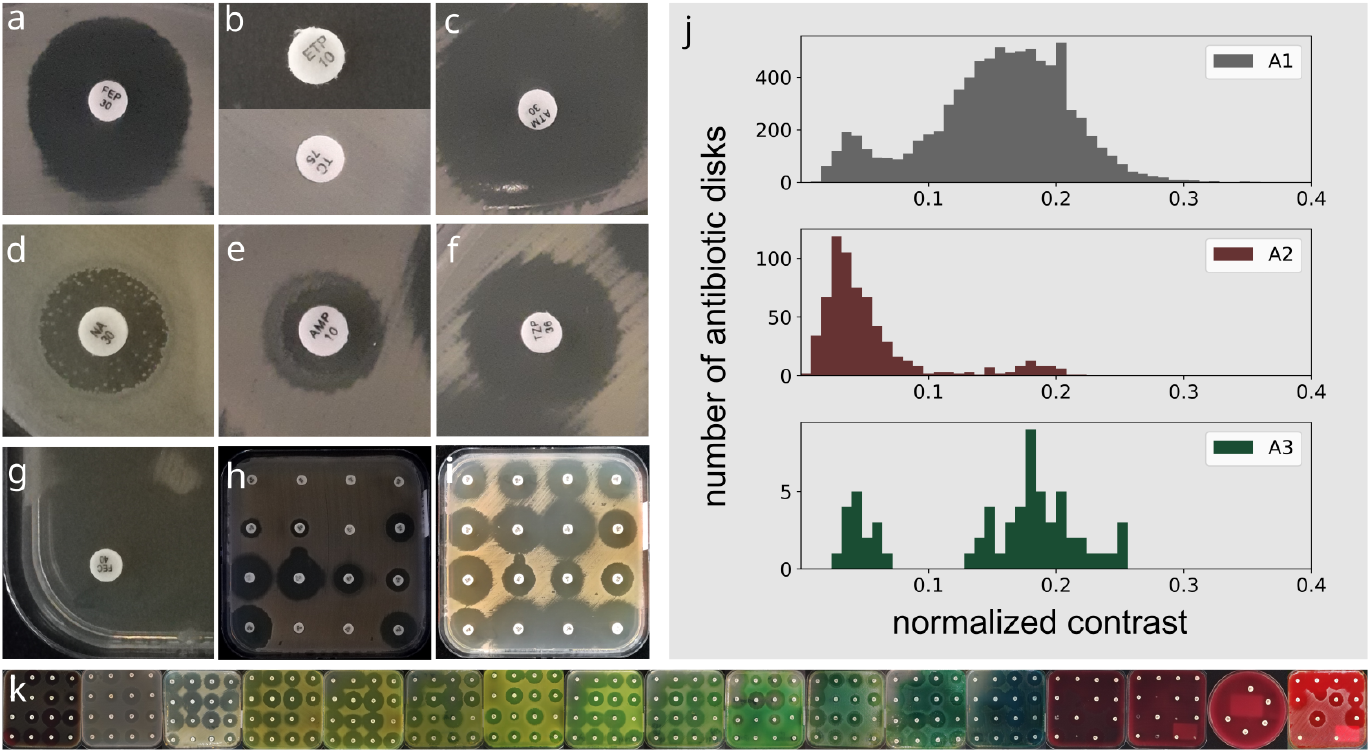
Variability in antibiogram pictures. Examples of difficult cases for diameter reading (a-h). Noncircular inhibition shape (a). Total or no inhibition (b). Light reflections (c). Colonies within the zone of inhibition (d). Double inhibition zones (e). Hazy borders (f). Inhibition zone overlap and plate borders (g). Low contrast (h) defined as the difference between the inhibition and bacteria intensity value, compared to a high contrast (i) image observed in A1. The histograms in (j) show the contrast variability observed in the benchmark data-sets (contrast is defined as the difference between the central gray levels of bacteria and inhibition, and normalized to the maximum available gray level). Observed variability in dominant hue (k).

The new algorithm for automatic diameter measurement, presented here, is referenced as SWITCH (Spatial Weighted Intensity Threshold CHangepoint). SWITCH operates a *k*-means clustering of the pixel intensity locally (around each antibiotic pellet) to classify inhibition and bacteria pixels (*k*=2). Successively, in order to find the inhibition zone boundary, it calculates and segments a radial profile *I*(*r*) measured in the surroundings of the antibiotic disk (up to the closest neighboring disk). For each value of *r* all pixels at distance *r* from the pellet center are considered. The value of *I*(*r*) is determined by the portion of pixels belonging to bacteria colonies (see details in Supplementary 7.4). Although SWITCH operates on a radial profile, the latter is calculated is a way that does not assume any preferential direction in the analysis of the image, which is important especially if the antibiotic disks are positioned by hand on the plate. Moreover it partially takes into account the texture of the colony, thereby increasing robustness to noise.

### 6.3 Susceptibility categorization and Interpretation

In this study, the susceptibility categorization of the tested antibiotics (S/I/R) is done by comparing the measured inhibition zone diameters to the EUCAST clinical breakpoints^7^. The breakpoint values are stored offline in the application, within the expert system knowledge base. This base, which contains also the expert rules and other expert system resources, is maintained and updated yearly by i2a^13,15^ (Montpelier, France).

In the context of AI, an Expert System is a program capable of taking reasoned conclusions from a given input, thereby simulating a human expert. Expert Systems have long been successfully used in microbiology^29^ and most commercial systems use them today. An Expert System consists of an “inference engine’ which takes reasoned conclusions on input information, based on a “set of rules” written by human experts.

The Expert System integrated in the App takes as input the diameter of the inhibition zones of the observed plate. It categorizes the susceptibility of the bacteria to the tested antibiotics and runs a coherence check and a final interpretation. The coherence check examines the input information and alerts the user if incoherent data are found (for example if an antibiotic is not coherent with the entered species, or if a natural resistance is not observed). The interpretation extrapolates the results classes of antibiotics and produces final alerts for important resistance mechanisms.

### 6.4 Auto-detection of resistance mechanisms

Certain resistance mechanisms to antibiotics can be detected by disk diffusion AST because they often produce inhibition zones with characteristic shapes^30,31^ (see Figure 7). These shapes appear between specific antibiotic disks. If the disks are close enough, the antibiotic molecules they diffuse can interact and produce a synergy effect against the bacteria or an induction of resistance. We used ML models to automatically recognize two particular shapes associated with two specific resistance mechanisms: synergy and induction, which can happen with *Extended-Spectrum β-lactamase (ESBL) production^32^* and *Macrolide-inducible resistance to Clindamycin* respectively.

**Figure 7:**
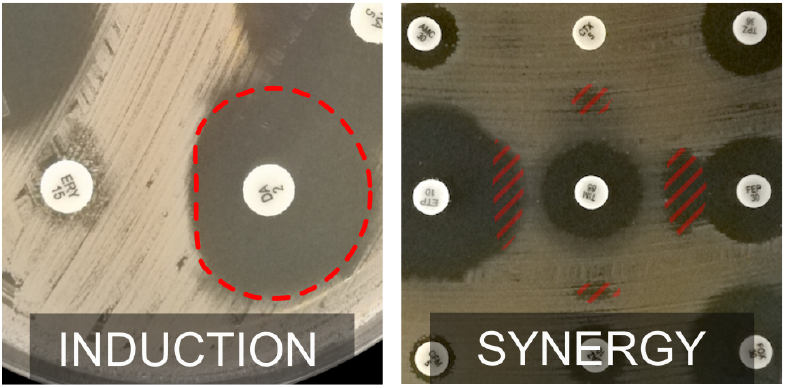
Characteristic shapes of inhibition zones due to resistance mechanisms.

For each of the two tests, we trained a neural network model to classify positive vs. negative images. The models to recognize Clindamycin-inducible resistance and ESBL reach an accuracy of 99.7% and 98%, respectively (see Supplementary 7.5 and 7.6 for details). However, the models have not been proven to perform well on a wide variety of images (varying in pellet arrangement, bacteria texture, etc). Also, since classification errors can have very serious consequences in AST interpretation and patient treatment, the App would need to ask the user for confirmation when automatically detecting a resistance mechanism. So, in the best case, including the resistance mechanism models in the App would bring only modest clinical improvements. Therefore, a ML-based automatic detection procedure was not included in the current version of the mobile application. Nevertheless, the App asks users if they see such shapes wherever they are likely to appear, and shows them examples for comparison (Figure 2).

## 7 Supplementary Information

### 7.1 Acquisition conditions

The precision of automatic measurements depends on image quality. In order to obtain the best possible results on images acquired with a smartphone camera, we designed a simple protocol and acquisition setup^IX^. First of all, the environmental illumination should be homogeneous and strong enough to comfortably read a printed text (e.g. the pictures should not be taken in a dark room nor close to a window or table lamp). The Petri dish should be placed uncovered on a flat, black, non-reflecting surface (e.g. a piece of black felt). A dark background enhances the bacteria/inhibition contrast and is a standard recommendation in the EUCAST guidelines for antibiogram reading^10^. Place on top of the Petri dish a sheet of black cardboard held by two objects of equal height. A hole the size of the phone camera is pierced in the middle of the cardboard and the pictures are taken through the hole with the phone lying on the cardboard. The distance between the Petri dish and the smartphone (i.e. the stand height) is determined by the camera’s field of view. It should be adjusted so that the Petri dish is well-fitted in the frame displayed by the camera.

The optics of smartphone cameras are not conceived for quantitative measurements, therefore small optical distortions are tolerated in production. However, initial exploration with standard calibration patters captured with several Android phones (Samsung A10, Huawei Y6, Pixel 3) suggested that optical distortion does not significantly impact measured diameters. In order for a smartphone camera to be suited for the AST measurements, the distortion it introduces should not impact the measurement more than the test sensibility (i.e. 1mm). In order to validate the smartphone camera, we generated and printed a fake antibiogram (see Figure 8). The ratio between the antibiotic disk and inhibition diameters in this picture is 25/6, so the App should measure a con-stant diameter of 25mm for each pellet. This simple test allows a rapid validation of smartphones cameras before using them for AST analysis.

**Figure 8:**
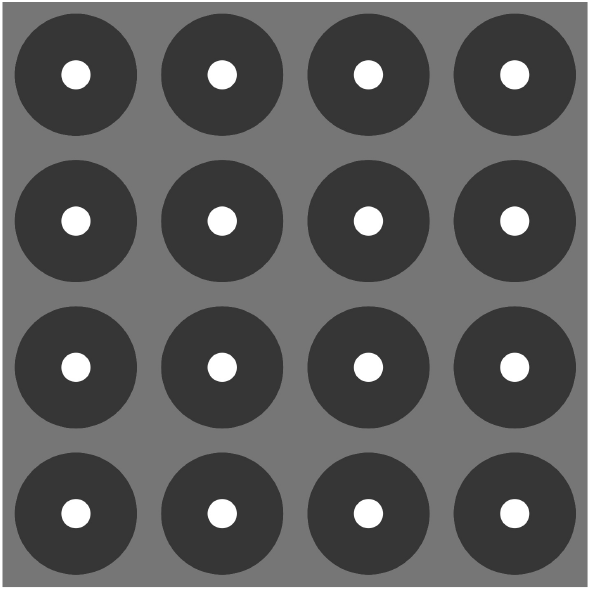
Numerically generated AST picture which can be printed and photographed to check if a smartphone camera is suited for analyzing antibiograms (right).

### 7.2 Pellet localization and extraction

If the antibiotic disks are placed on the antibiogram with a dispenser, their relative position is known in advance and can be exploited^20^. Unfortunately, dispensers are not always available and placing the antibiotic disks by hand remains a common practice in many laboratories, especially in resource-limited settings. Therefore, we decided not to rely on any assumptions regarding pellet position.

The detection procedure is based on the hypothesis that the pellets are white disks: their shape is approximately circular, and the intensity levels of the associated pixels in the digitized image are high. The blue channel is taken from the cropped image of the Petri dish: we observed that in pellets usually have higher contrast from the rest of the image in the blue channel. Contrast is enhanced by histogram normalization, then an intensity threshold is applied to distinguish pellets from non-pellet pixels. The threshold is 0.97 times the maximum allowed intensity level (white).

The image is de-noised with standard mathematical morphology operations. Then, the contours of the connected components are extracted ^33^ and filtered according to their relative size in the image and to their thinness t:

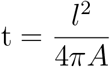

where *l* is the length of the contour and *A* is the enclosed area. By construction, t ≃ 1 for a contour close in shape to a circle.

This eventually allows to discard bright objects such as light speckles. The contours are considered to belong to antibiotic disks if their diameter is within 1/30 and 1/4 of the image largest size and th < 1.05.

If an antibiotic pellet is not detected automatically, the user needs to add it manually in order to continue the analysis. The detection precision influences the further analysis results. For this task we implemented a semi-automatic procedure that takes as input the approximate coordinates of the pellet center (e.g. the user clicks on the pellet) and then accurately finds the pellet center and diameter with a Hough circle transform ^34^. The previously collected information of average pellet size in the image is used to filter in the transform space.

### 7.3 Antibiotic disk label reading by ML

Antibiotic name recognition is treated as a multi-class problem because there are as many codes to be recognized as there are antibiotics. This supervised learning problem is tackled via a con-volutional neural network (CNN) trained using 18000 images of antibiotic disks captured with a smartphone camera. More precisely, the images were split in two sets: a training set of 14000 images and a test set of 4000 images used also for validation.

The set contains 65 different labels corresponding to the pellets of two manufacturers (Liofilchem and i2a). We estimate that across the major disks manufacturers, there are over ≃ 200 different labels. Our model should eventually be able to identify all of those labels.

The input images are first converted to gray-scale and resized to 64×64 pixels. Then, the intensity value of each pixel is normalized by subtracting the mean intensity and dividing by the intensity standard deviation. The model takes a standardized pellet image as input and returns an array of 65 values between 0 and 1 normalized to 1. These values are interpreted as the probability of belonging to a specific class (i.e. label for a given antibiotic).

The model is a CNN with three convolution layers described in Table 3. Each convolution layer is followed by max-pooling with a down-scaling factor of 2. The classification output is obtained with a random 50% dropout and a final dense layer of 65 units. We used cross-entropy as loss function.

**Table 3:** Convolution layers in the CNN model used to classify the antibiotic disk labels

| layer type | kernel size | output size | activation |
| --- | --- | --- | --- |
| 2D convolution | 8 | 32 | relu |
| 2D convolution | 4 | 64 | relu |
| 2D convolution | 2 | 64 | relu |

The model is evaluated primarily on its accuracy. Since the data set doesn’t cover all the antibiotic codes that could be encountered in a real use case, a second metric was introduced to evaluate the model behavior on unknown labels (hereafter “outliers”). To simulate this behavior, 5 classes (each class corresponds to a label) are held out from the dataset and the model is trained on the remaining 60. After defining a threshold on the model output, we evaluate:

- for the 60 classes from the test data

– the accuracy on the predictions above the threshold (“accuracy on known labels”)
– the percentage of samples that fell below the threshold (“false negative rate on known labels”)
- for the 5 classes held out and taken from the test data

– the percentage of samples that fell above the threshold (“false positive rate on outliers”)

A single model showed good accuracy, but a simple threshold on the output values was not good enough to minimize classification errors of outliers. Therefore, we developed an ensemble model with an entropy threshold to achieve a better trade-off between accuracy and outlier classification error. We trained ten identical instances of this model with same hyper-parameters, same training data, but different random initialization. The models were trained using Tensorflow^23^ in Python.

Classification is obtained by feeding the same pre-processed antibiotic disk image to each model of the ensemble, the ten classification outputs are averaged, and the information entropy of the average is calculated. If the entropy is larger than a fixed threshold, the *argmax* of the output average determines which classes the label belongs to. Otherwise, the label is considered as unknown. The threshold is chosen to optimize the trade-off between optimizing accuracy and minimizing out-of-distribution error. From the user point of view, if a disk is classified as unknown, the users must provide the label based on visual inspection. The list of possible labels are sorted according to their output prediction value, which usually brings to the top the best candidates in case of false negatives (known pellets classified as unknown).

The ensemble model achieves 99.97% accuracy on the test set. Using an ensemble of 10 models and an entropy threshold, we achieved: 100% in-distribution accuracy, 5% in-distribution false negative rate and 1% out-of-distribution false positive rate.

Additionally, the pictures of antibiogram set A1 can also be considered as a true test set because the model never saw them during training. The model reaches 100% accuracy on this test set, as discussed in Results.

### 7.4 Inhibition disk diameter measurement

Diameter measurement consists of three steps: image pre-processing, intensity radial profile extraction and segmentation.

#### 7.4.1 Pre-processing

The growth medium of the Petri dish is automatically determined based on the saturation, hue and intensity of the Petri dish image. First, we calculate the mean saturation, hue and intensity over the 50% most saturated pixels in the image, then we use these values to classify the growth medium. The classifier is built with a logistic regression on more than 600 labeled images (see Figure 9) and achieves 100% accuracy on this data set. If a red growth medium (HM-F) is detected, we use only the blue and green channels of the image when converting it to gray scale, in order to enhance the output contrast. Otherwise all three channels are used.

**Figure 9:**
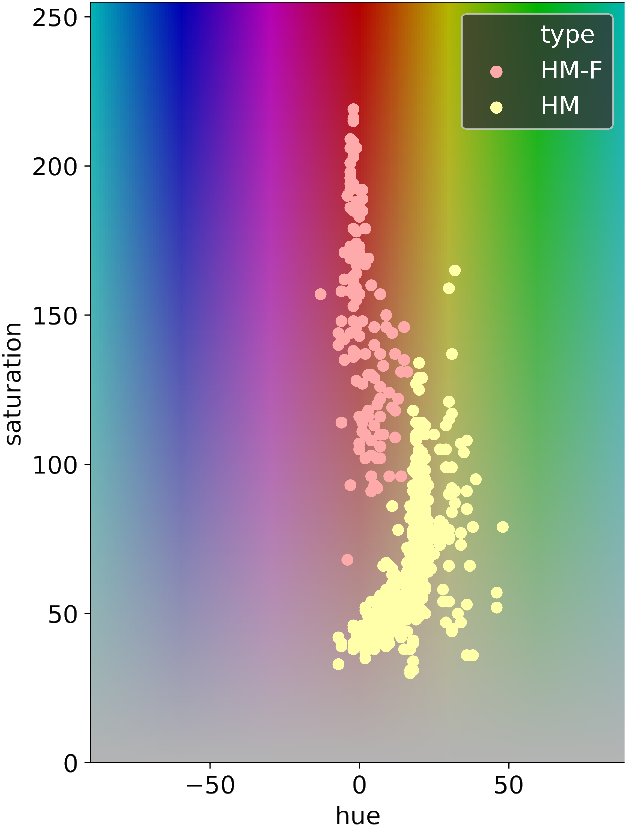
distribution of the mean HUE and saturation values of the AST pictures in data sets A1 and A2. Each point represents one image.

**Figure 10:**
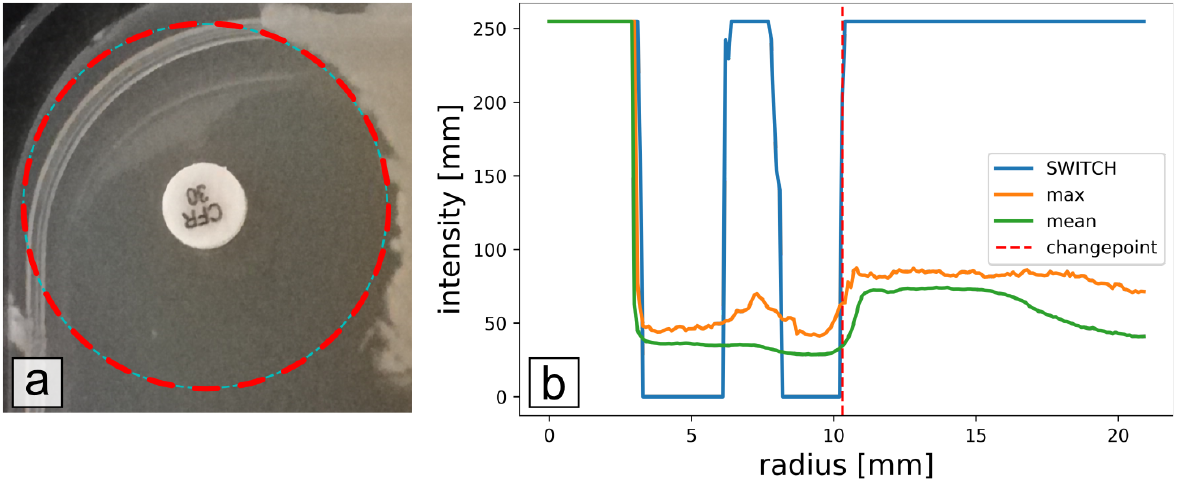
Radial intensity profile extraction. The region of interest of an antibiotic pellet is displayed in (a). Different radial profiles are plotted in (b) by taking the max or mean intensity value at distance *r*. The blue profile is calculated by the SWITCH algorithm proposed here. The peak of the signal in the inhibition zone is due to light reflection on the agar, but the segmentation correctly ignores it. In the mean profile, since the borders of the inhibition zone are not flat, the signal ramps up smoothly, introducing uncertainty in the change-point location, then the signal drops down because of inhibition zone overlap. These problems do not affect SWITCH.

Successively the image is pre-processed for the diameter measurement. The Petri dish image is down-scaled to a fixed resolution of 10 pixel/mm, which provides a reasonable trade off between precision and measurement speed. Then, all the pixels belonging to the plate borders are masked. This eliminates disturbing light reflections (due to surface tension, the agar is not flat close to the plate’s borders). The plate image is segmented in Regions Of Interest (ROIs), rectangular subimages centered on each antibiotic disk. Each ROI includes the area surrounding the disk up to the first-neighboring disks.

For the whole image first and then for each region of interest, a *k*-means classification of the pixel intensity is performed with *k*=2 (inhibition and bacteria) and the center values are stored. Since the *k*-means classification with *k*=2 is not optimal for very large inhibition zones or zones with no inhibition at all, in this case the *k*-means centers calculated on the whole image are used.

#### 7.4.2 Radial profiles

The intensity of the pixels around the antibiotic disk is observed for each ROI in order to calculate a radial intensity profile *I* (*r*). The profile extraction procedure takes into account both the intensity and the amount of bacteria at a given *r*. Without considering any privileged direction, all pixels at a given distance r from the antibiotic disk center are considered. Then *I* (*r*) is given a score between 0 and 1 depending on the number *n_b_* of “bacteria pixels”. “Bacteria pixels” are defined as those pixels which have an intensity above the threshold *I_th_*. The bacteria intensity threshold is calculated as:

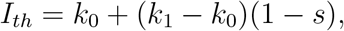

where *k*_0_ and *k*_1_ are the local *k*-means centers for inhibition and bacteria pixel intensities. *s* is defined as the reading sensibility. It can be fixed between 0 and 1 or automatically determined based on the image local and global contrast *c_l_*. Contrast is defined as (*c* = *k*_1_ − *k*_0_), local contrast is based on local *k*-means (over the antibiotic disk ROI) and global contrast on global *k*-means (whole image).

If automatically determined, s is equal to 0.05 for very low contrast images (contrast≤25) where the signal/noise ratio is low, otherwise

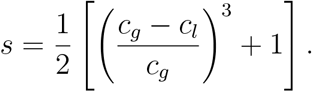

If the local and global contrast values are similar s ≃ 1/2. Otherwise, it is adapted according to the ratio *c_l_/c_g_* in order to compensate noise and local intensity variability in the image.

The critical number of bacteria pixels *n_b_* represents a spatial threshold that allows detecting the presence of bacteria at distance *r* even if the shape of the inhibition zone is not circular (e.g. inhibition overlap and plate borders). We choose *n_b_* = 1mm in the image scale.

Finally, *I*(*r*) is segmented using a least-square fit of a step function with one degree of freedom: the change-point, which is interpreted as the inhibition zone radius in pixels. A maximum inhibition diameter can be specified by the user, in this case the reported diameters would be capped by that maximum (40mm by default).

The absolute diameter difference between automatic measurements and control on the test antibiotics sets are reported in Table 4. The larger absolute diameter difference in A1 and A2 can be attributed to the fact that the control diameter (measured by a commercial automatic system) was not systematically adjusted if adjustment did not have an effect on the categorization.

**Table 4:** Absolute diameter difference (a.a.d.) between automatic measurement and control.

| AST set | control | images | antibiotics | a.d.d. quantiles [mm] |  |  |
| --- | --- | --- | --- | --- | --- | --- |
|  |  |  |  | 25% | 50% | 75% |
| A1 |  |  |  |  |  |  |
| overall | SIRscan | 572 | 8168 | 0.8 | 1.8 | 3.0 |
| standard | SIRscan | 561 | 8042 | 0.8 | 1.8 | 3.0 |
| problematic | SIRscan | 9 | 126 | 2.4 | 6.3 | 20.8 |
| A2 |  |  |  |  |  |  |
| overall | SIRscan | 75 | 649 | 0.6 | 1.4 | 2.8 |
| standard | SIRscan | 73 | 620 | 0.6 | 1.4 | 2.6 |
| problematic | SIRscan | 2 | 29 | 11.0 | 17.0 | 22.0 |
| A3 |  |  |  |  |  |  |
| overall | manual | 64 | 784 | 0.0 | 0.8 | 1.4 |
| overall | manual av. | 64 | 98 | 0.1 | 0.6 | 1.3 |

**Table 5:**
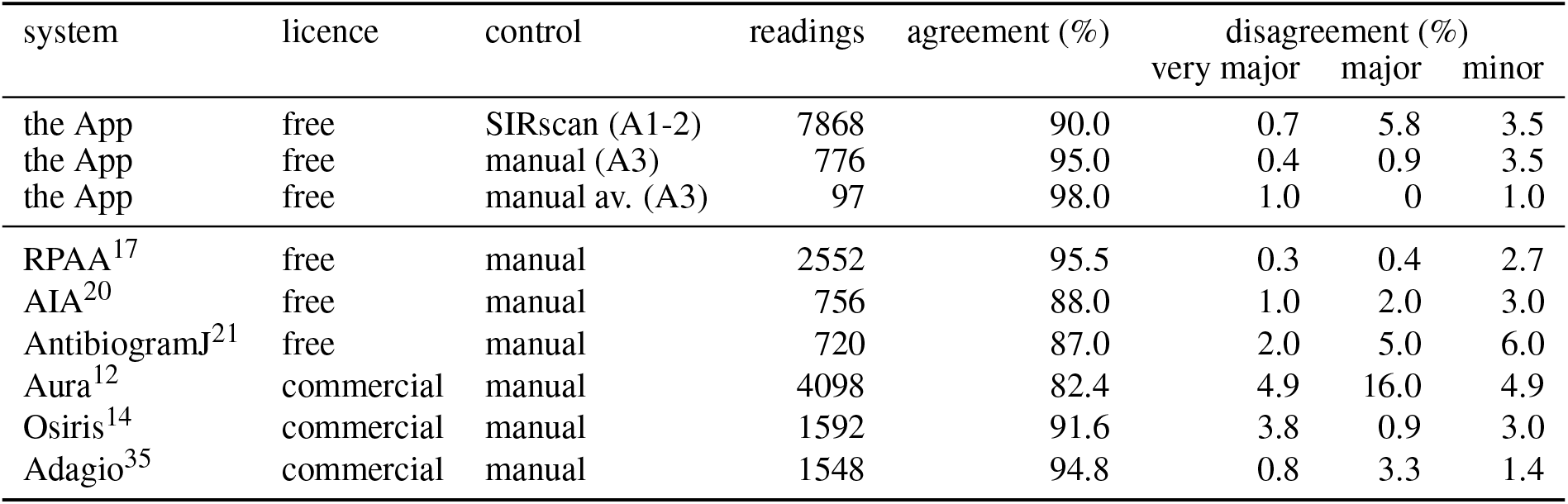
Comparison of categorization agreement/disagreement of the App’s performance shown in this paper versus similar studies concerning other automatic systems. This comparison should be taken with caution because the datasets used in these studies are different under many aspects (e.g. species distribution, manual measurement procedure, image acquisition and outliers dropout).

| system | licence | control | readings | agreement (%) | disagreement (%) |  |  |
| --- | --- | --- | --- | --- | --- | --- | --- |
|  |  |  |  |  | very major | major | minor |
| the App | free | SIRscan (A1-2) | 7868 | 90.0 | 0.7 | 5.8 | 3.5 |
| the App | free | manual (A3) | 776 | 95.0 | 0.4 | 0.9 | 3.5 |
| the App | free | manual av. (A3) | 97 | 98.0 | 1.0 | 0 | 1.0 |
| RPAA<sup>17</sup> | free | manual | 2552 | 95.5 | 0.3 | 0.4 | 2.7 |
| AIA<sup>20</sup> | free | manual | 756 | 88.0 | 1.0 | 2.0 | 3.0 |
| AntibiogramJ<sup>21</sup> | free | manual | 720 | 87.0 | 2.0 | 5.0 | 6.0 |
| Aura<sup>12</sup> | commercial | manual | 4098 | 82.4 | 4.9 | 16.0 | 4.9 |
| Osiris<sup>14</sup> | commercial | manual | 1592 | 91.6 | 3.8 | 0.9 | 3.0 |
| Adagio<sup>35</sup> | commercial | manual | 1548 | 94.8 | 0.8 | 3.3 | 1.4 |

**Table 6:** Effect of resolution degradation on dataset A3 (averaged). The results are substantially the same up to resolution = 3 Megapixel. At resolution = 1 megapixel, 6 antibiotic disks are missed out of the total 98, but the classification error remains constant.

| resolution [megapixel] | missed atb disks | unread labels | a.d.d. quantiles [mm] |  |  | disagreement |  |  |
| --- | --- | --- | --- | --- | --- | --- | --- | --- |
|  |  |  | 25% | 50% | 75% | very major | major | minor |
| original (12.8) | 0 | 0 | 0.12 | 0.60 | 1.27 | 1 | 0 | 1 |
| 9.0 | 0 | 0 | 0.13 | 0.55 | 1.25 | 1 | 0 | 1 |
| 6.0 | 0 | 1 | 0.15 | 0.53 | 1.25 | 1 | 0 | 1 |
| 3.0 | 0 | 0 | 0.15 | 0.57 | 1.19 | 1 | 0 | 1 |
| 1.0 | 6 | 2 | 0.31 | 0.53 | 1.18 | 1 | 0 | 1 |

#### 7.4.3 Published algorithms for measuring diameters

Among the different algorithms for this specific task we could find in the literature^17,18^ do not prove to be resilient to the common problems of non-homogeneous or non-circular inhibition zones. Same for ^19^, which takes into account both intensity and texture with a Student’s t-test, but is sensitive to noise and assumes that the inhibition and bacteria have homogeneous textures. A recent work^20^ calculates a threshold for segmentation only along 4 segments centered on the antibiotic pellet, with the risk of losing information.

### 7.5 Resistance mechanisms: D-shape (MLSb inducible resistant Staphylococcus aureus)

The raw input to the training pipeline consists of photos of entire Petri dishes used for AST. Photos were taken using a smartphone camera at an MSF field hospital in Amman, Jordan during the course of regular AST processing. As a result, the photos represent similar conditions to those under which the App will eventually be used. Only relevant examples (those containing adjacent pairs of Clindamycin and Erythromycin pellets) are considered. The training data includes 69 positive and 143 negative examples, each image is labeled as “D-Zone or “not D-Zone.

First, the photos are preprocessed. The first four preprocessing steps aim to standardize input characteristics across examples (see Figure 12):

1. Identify the Clindamycin and Erythromycin pellets using the pellet label recognition strategy described above.
2. Rotate the image such that Erythromycin and Clindamycin are aligned along the x-axis, with Erythromycin on the left.
3. Crop out the biggest region surrounding Clindamycin that does not include other pellets.
4. Overlay the Clindamycin pellet with a 6 mm white circle in order to mask the printed text, which has no interest in the classification problem. In the interest of a small, lightweight model, the final two steps simply reduce input size without loss of model accuracy. As a result, the numerical input to the machine learning model is simply a 32×32 integer matrix, where each entry is a value between 0 (black) and 255 (white).
5. Convert the image from color to gray-scale
6. Downsize the image to 32 pixel × 32 pixel.

**Figure 12:**
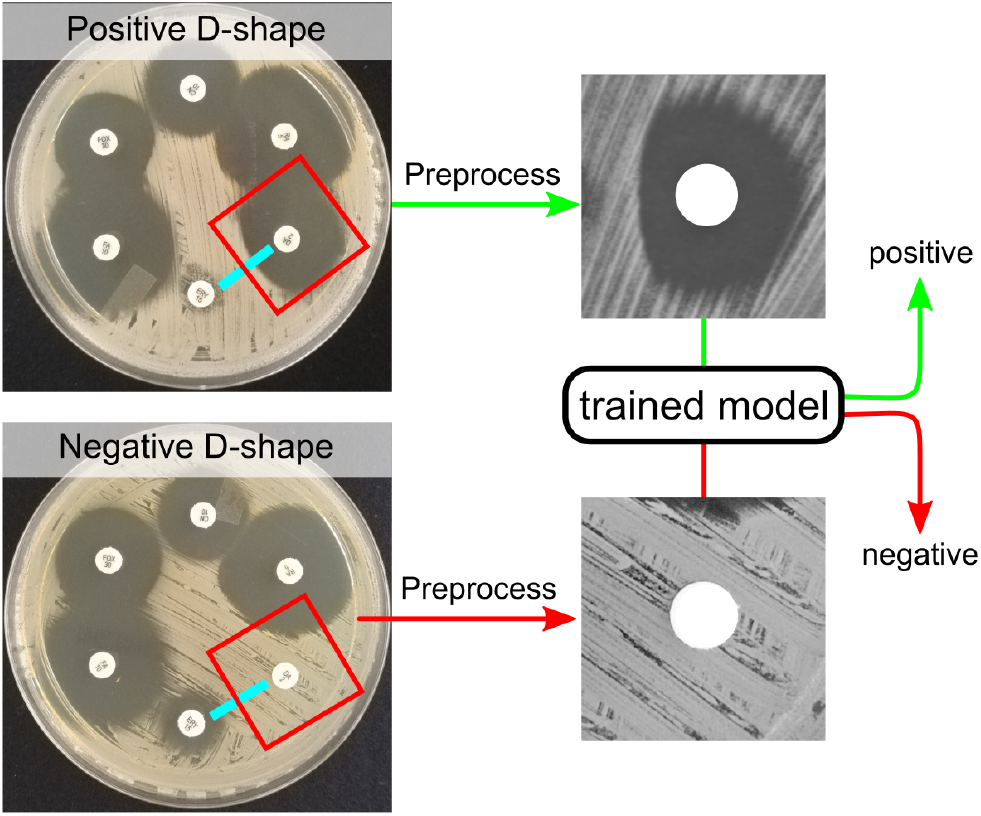
D-Zone prediction workflow.

We report the model’s train and test accuracy, i.e. the proportion of examples for which the model correctly predicts the known label. Since the data-set is small, a 70/30 split results in 148 training samples and 64 test samples, therefore we evaluate model structures by retraining the same model structure on multiple different train/test splits. However, the model is never tested on the same images as it was trained. This approach allows us to compare different model architectures, whereas a single train/test split would result in ties.

Models were trained using TensorFlow in Python. After experimenting with multiple neural network architectures, the architecture with the best performance has the following characteristics:

- Input size of 32 pixel × 32 pixel,
- 73,309 total trainable parameters,
- Binary cross-entropy loss function
- Adam optimization,
- input size of 32 pixel × 32 pixel,
- layers: 3x 2-D convolution layers rectified linear unit activation, with 32, 64, and 128-dimensional outputs respectively,
- Dropout layer to avoid over-fitting,
- Densely-connected layer of size 50 with sigmoid activation (for binary classification).

Across 30 randomly-chosen train/test splits, the best model achieves on average 100% training accuracy and 99.74% test accuracy. Other metrics give similar results (F1 99.65% and AUC 1.0), as expected given the high accuracy score.

D-Zone detection is a relatively easy ML problem, obtaining 99.74% test accuracy with only 212 training images. The D-Zone model has not been evaluated on out-of-distribution images.

### 7.6 Resistance mechanisms: ESBL

For the ESBL problem, we have access to two distinct data-sets (see Table 7. Each data-set consists of images of AST plates. Each image contains pellets arranged to perform a Double Disk Synergy test and each is labeled as “ESBL” or “not ESBL”. The two data-sets differ highly in different aspects such as image quality and intensity contrast, bacteria culture texture, pellet arrangement, specific antibiotics used.

**Table 7:** Training data-sets for the synergy model.

| Data-set name | D1 | D2 |
| --- | --- | --- |
| acquisition device | smartphone camera | SIRscan system |
| Number of ESBL positive examples | 241 | 1344 |
| Number of ESBL negative examples | 181 | 818 |
| Center pellet | Amoxicillin/Clavulanic acid 30 μg | Amoxicillin/Clavulanic acid 30 μg or Ticarillin/Clavulanic acid 75+10 μg |

**Table 8:** ESBL model accuracy results.

| train set (%) |  | test set (%) |  | preprocessing | accuracy (%) |  | F1 | AUC |
| --- | --- | --- | --- | --- | --- | --- | --- | --- |
| D1 | D2 | D1 | D2 |  | train | test |  |  |
| 70 | - | 30 | - | light | 99.83 | 98.04 | 98.89 | 99.84 |
| - | 70 | - | 30 | light | 98.89 | 97.98 | 99.06 | 99.70 |
| 100 | - | - | 100 | light | 99.93 | 66.77 | 76.90 | 67.17 |
| - | 100 | 100 | - | light | 99.75 | 69.23 | 62.09 | 64.75 |
| 100 | - | - | 100 | heavy | 99.97 | 58.96 | NA | NA |
| - | 100 | 100 | - | heavy | 99.59 | 59.16 | NA | NA |

Before training a neural network on the images, we preprocess them to standardize across images and extract relevant regions:

1. Crop out a 35mm region surrounding amoxicillin-clavulanate. In both our data-sets, 35mm is sufficiently large to encompass amoxicillin-clavulanate and the surrounding 3rd-generation cephalosporins.
2. Convert the image to gray scale.
3. Normalize image contrast such that the intensity threshold is similar across all images.
4. Overlay the amoxicillin-clavulanate pellet with a pure black circle.

Alternative preprocessing approaches did not introduce significant improvement of the prediction accuracy. For example we tested: 1. blur the image to eliminate unnecessary details such as bacteria streaks or 2. classify each pixel as bacteria vs. inhibition using an intensity threshold computed by *k*-means clustering (we will refer to this strategy as “heavy preprocessing” in this section).

As in the case of D-Zone, train and test accuracy is reported. However, since in the ESBL case we have access to two distinct data-sets, we also examine model transferability. Therefore, we examine train and test accuracy within each of the following setups:

- Train and test on disjoint sets of images drawn from the same data-set
- Train and test on disjoint sets of images, where both the test and train sets are drawn from the combination of both data-sets
- Train on one data-set and test on the other

The model is over 97% accurate when its test set is drawn from the same distribution as its training set. However, when the training and test sets are of different origin, the model performs no better than random. Out-of-distribution examples are of interest because we cannot control the quality of images that the app will eventually be used to analyze. Specifically to address this problem, we tested the “heavy” preprocessing approach but did not improve cross-data-set accuracy. As more training examples are added, model accuracy increases only until 300 training examples are used. Additionally, augmenting the training data-set by transforming (e.g., rotating) images did not improve validation accuracy. Therefore, obtaining more similar data is not expected to improve results. The results obtained with other metrics (F1 and ROC AUC) justified by the unbalanced data classes, confirm good performance for intra-dataset models and a much lower performance for inter-dataset models.

ESBL classification is a much more difficult machine learning problem than D-Zone classification (see Supplementary 7.5). This difficulty might be due to the very large variability in the ESBL-positive examples (see Figure 13). While D-Zone examples are all very similar to each other, ESBL-positive examples may show different shapes, depending on the relative position and distance of the involved antibiotic disks, on the texture of the bacteria and on the quality of the picture.

**Figure 13:**
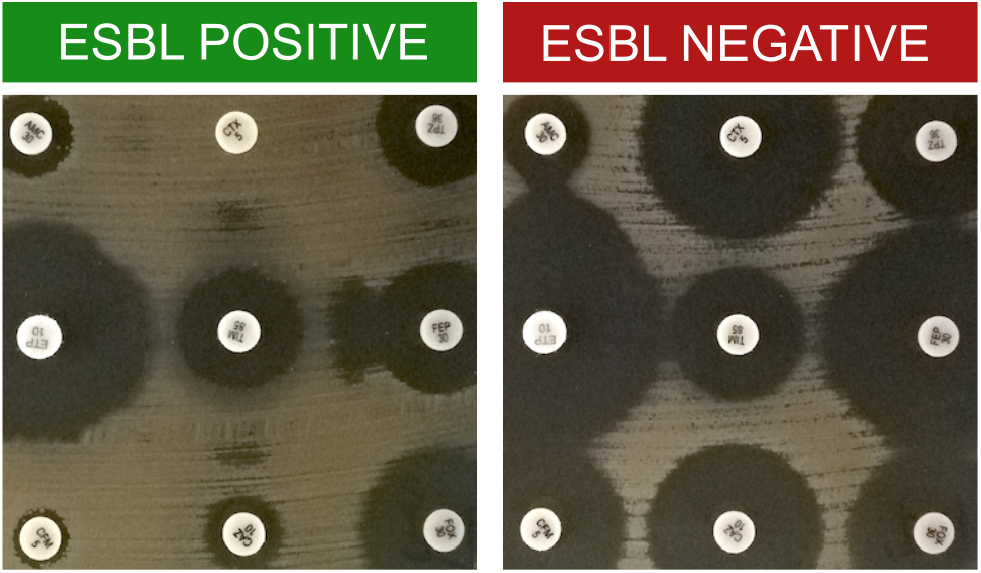
Examples of ESBL positive and negative images. The shapes of the inhibition zone can be very different.

## 8 Acknowledgments

The App is supported by: the Médecins Sans Frontières Foundation, the Laboratoire de Mathématiques et Modélisation d’Évry, the Génoscope, the bacteriology laoratory of the Universitary Hospital Henri Mondor, Créteil France. For its potential impact, this project was awarded the 2019 Google AI Impact Challenge^36^, selected among over 2600 applicants worldwide.

We are expecially thankful to Dominique Boissinot and Alain Jean of i2a for their fundamental help in the development of the Expert System. We Thank all the organizers of the Google AI Impact Challenge and the people of Google.org, among others Anna Achilles, Sebastien Floodpage and Mollie Javerbaum for their invaluable help.

## 9 Author Contributions

- **Marco Pascucci** designed and coded the Image Processing system (including the novel algorithms) except for the ML based label recognition part. He did the bibliographical research and wrote the largest contribution to the manuscript. He did a feasibility study and built the first prototype of the application (which entered the Google AI impact challenge). He studied, evaluated, and prepared the knowledge base of i2a’s Expert System to be used within the app, which motivated the adoption of this component. He conceived the first global design of the app’s architecture. He designed the experiments concerning AST sets A1 and A2. Finally He did all the analysis illustrated in the paper and interpreted the results.
- **Guilhem Royer** designed the experimental procedure for AST set A1 and A2 and collected the data. He analyzed the data and wrote the paper. As expert clinical microbiologist, he gave major contributions in understanding the needs of the development, concerning both the image processing and the expert system.
- **Jakub Adamek** developed the mobile application, especially the UX, the integration of Tensorflow and tests.
- **David Aristizabal** developed the mobile application. He did a very careful review of the paper manuscript.
- **Amine Bezzarga** Developed the expert system engine which uses the knowledge base of i2a, embedded in the application. He contributed to other aspect of the mobile app development and deployment.
- **Laetitia Blanche** was product Manager of the application since august 2019, she worked on the definition of the user journeys, the needs specification and coordination of the development and release of the mobile application.
- **Guillaume Boniface-Chang** was product Manager on the Google side. He built and trained the model that recognises the antibiotic pellets images, with the help of Gabriel Dulac-Arnold. The sections of the paper describing this features are largely taken from the documentation he wrote.
- **Alex Brunner** was a UX researcher in the first google fellows cohort, contributed in understanding users needs and designing the user journeys.
- **Philippe Cavalier** was consultant microbiologist and contributed to the design of the experimental protocol concerning AST set A3.
- **Christian Curel**, the CEO of i2a, who provided the Expert System knowledge base.
- **Gabriel Dulac-Arnold** helped in the different aspect of the app concerning ML.
- **Nada Malou** provided expert insights that proved to be critical for the overall definition of the project throughout all phases of development: mapping user journeys and pain points, identifying and prioritizing strategic requirements for the app, and enabling access to on-field testing. She designed the experimental protocol concerning AST set A3. This protocol defined objectives and key metrics, the workflow to collect them as well as underlying requirements. She also led on the submission of this protocol to MSF Ethical Review Board. She coordinated training and recruitment for the research assistants that performed the measurements in AST set A3. She tested the Expert System that was implemented as a core component of the app, by providing a representative set of complex interpretation cases.
- **Clara Nordon** as director of the MSF Foundation took care that all the necessary conditions were reunited in order to ensure the conception and development of the app.
- **Vincent Runge** gave important inputs and contributions to the automatic measurement of the inhibition diameters.
- **Franck Samson** gave determinant contributions in developing the first prototype of the UI of the mobile app;
- **Ellen Sebastian** contributed to the mobile application development. Ellen worked with Guillaume Boniface-Chang, Gabriel Dulac-Arnold and M. Pascucci on the proof of principle of the automatic recognition of the resistance mechanisms by classification of the shape of the inhibition zones, for which she was the main contributor. The sections of the paper describing these features are largely taken from the documentation she wrote. She did accurate reviews of the whole paper.
- **Dena Soukieh** was a UX designer for the app in the first cohort of Google fellows, she contributed in designing the UX.
- **Jean-Philippe Vert** provided helpful discussions about ML, discussed the results and reviewed the manuscript.
- **Christophe Ambroise** designed and directed the study, interpreted the results and wrote the paper.
- **Mohammed-Amin Madoui** had the original idea of the application, designed, interpreted the results, directed the study and wrote the paper.

## Footnotes

I https://fondation.msf.fr/en/projects/antibiogo

II available at https://mpascucci.github.io/AST-image-processing

III https://youtu.be/0hNr9zTu6ig

IV G.L.A.S.S. https://www.who.int/glass/en

V WHOnet https://whonet.org

VI AST image processing: https://github.com/mpascucci/AST-image-processing

VII App access form: https://form.typeform.com/to/qEGVBzbu

VIII https://mpascucci.github.io/AST-image-processing

IX https://mpascucci.github.io/ASTapp-protocol

